# The Na_V_1.5 auxiliary subunit FGF13 modulates channels by regulating membrane cholesterol independent of channel binding

**DOI:** 10.1101/2025.03.19.644183

**Authors:** Aravind Gade, Mattia Malvezzi, Lala Tanmoy Das, Maiko Matsui, Cheng-I J. Ma, Keon Mazdisnian, Steven O. Marx, Frederick R. Maxfield, Geoffrey S. Pitt

## Abstract

Fibroblast growth factor homologous factors (FHFs) bind to the cytoplasmic carboxy terminus of voltage-gated sodium channels (VGSCs) and modulate channel function. Variants in FHFs or VGSCs perturbing that bimolecular interaction are associated with arrhythmias. Like some channel auxiliary subunits, FHFs exert additional cellular regulatory roles, but whether these alternative roles affect VGSC regulation is unknown. Using a separation-of-function strategy, we show that a structurally guided, binding incompetent mutant FGF13 (the major FHF in mouse heart), confers complete regulation of VGSC steady-state inactivation (SSI), the canonical effect of FHFs. In cardiomyocytes isolated from *Fgf13* knockout mice, expression of the mutant FGF13 completely restores wild-type regulation of SSI. FGF13 regulation of SSI derives from effects on local accessible membrane cholesterol, which is unexpectedly polarized and concentrated in cardiomyocytes at the intercalated disc (ID) where most VGSCs localize. *Fgf13* knockout eliminates the polarized cholesterol distribution and causes loss of VGSCs from the ID. Moreover, we show that the previously described FGF13-dependent stabilization of VGSC currents at elevated temperatures depends on the cholesterol mechanism. These results provide new insights into how FHFs affect VGSCs and alter the canonical model by which channel auxiliary exert influence.

## Introduction

While pore-forming subunits of ion channels are responsible for ion conduction, auxiliary subunits critically influence channel function, as reflected by variants in multiple auxiliary subunit genes that are associated with arrhythmia syndromes (1, 2). Among their roles, auxiliary subunits control trafficking of channels to the sarcolemma, target channels to specific subcellular locations on the membrane, and tune biophysical properties of the channels inserted into the membrane. Auxiliary subunits are thought to affect channel function via direct binding mechanisms. This direct interaction model, however, is incomplete as some ion channel auxiliary subunits only “moonlight” as channel auxiliary subunits while serving other essential roles independent of channel interaction. A prime example is calmodulin, the most abundant intracellular Ca^2+^ binding protein and a regulator of numerous Ca^2+^ signaling events, that also binds directly to, and serves as an auxiliary subunit for, multiple channels such as voltage-gated sodium channels (**VGSCs**), Ca^2+^ channels, and K^+^ channels (3–12). Whether the other cellular functions controlled by these auxiliary subunits affect channel function, however, is underexplored.

Here, we focus on the fibroblast growth factor homologous factor (**FHF**) subfamily of fibroblast growth factors (**FGFs**). The FHFs (FGF11-FGF14) comprise a noncanonical FGF subset that is not secreted (13, 14) and does not function as growth factors (15). FHFs can bind to the cytoplasmic VGSC C-termini (16) and modulate various aspects of VGSC function, one of the most prominent of which is increasing channel availability (i.e., a depolarizing shift in the V_1/2_ of steady-state inactivation [**SSI**]) (17–19). Among the FHFs, human cardiomyocytes express predominantly *FGF12* and to a lesser extent *FGF13* (19–21). Variants of *FGF12* have been associated with inherited ventricular arrhythmias (20, 22) including Brugada syndrome (**BrS**), an arrhythmia characterized by reduced Na_V_1.5 current. A genome-wide association study (**GWAS**) identified *FGF13* as a risk locus for atrial fibrillation (23) (**AF**) and postoperative AF is associated with reduced *FGF13* expression (24). Rodents predominantly express *Fgf13* in the ventricle (19) and cardiomyocyte-restricted *Fgf13* knockdown or knockout (**KO**) affects VGSC currents (19, 25–28). The most consistent observed effect in *Fgf13* KO mice—present in all studies to date (19, 25–28)—is a decrease in channel availability. Further, in *Fgf13* KO cardiomyocytes elevated temperature slows conduction velocity through the ventricular myocardium (26), a feature associated with BrS, in which affected individuals are at ∼20-fold increased risk of a type I BrS ECG pattern and arrhythmias in the setting of a fever (29). How FGF13 protects against the effects of elevated temperature in the myocardium or how reduced FGF13 contributes to postoperative AF in humans is not known, thus preventing development of targeted therapies.

The working hypothesis for how FHFs regulate VGSCs is by a structurally defined direct interaction between the FHF and the cytoplasmic VGSC C-terminus via binding determinants within FHFs and VGSCs that are conserved among their respective family members (30, 31). A mutation within the cardiac VGSC Na_V_1.5 C-terminus that inhibits FHF binding is associated with ventricular arrhythmias and sudden cardiac death in a five-generation family (32), underscoring the importance of this interaction. Similarly, a mutation perturbing the corresponding interaction site on FGF12, a FHF expressed in human brain and heart, affects neuronal VGSC function (33) and is associated with a seizure disorder (34); and also affects cardiac VGSC channel currents when the mutant FGF12 is expressed in rodent cardiomyocytes (33). A separate variant in *FGF12* that impairs FGF12 interaction with Na_V_1.5 has been associated with BrS (20). As evolutionarily derived from FGFs, FHFs likely have multiple effects beyond direct binding to VGSCs; and these other roles could affect VGSCs via a binding independent manner. Indeed, FGF13 stabilizes microtubules (14, 35, 36), which are critical for trafficking and targeting VGSCs (37).

Here, using a structurally informed FGF13 mutant unable to bind VGSCs, we show that FGF13 exerts critical components of its VGSC regulatory functions, including the canonical shift in SSI, via binding-independent mechanisms.

## Results

We employed a constitutive cardiac-specific knockout model (c*Fgf13^KO^*) to investigate how FGF13 regulates VGSCs in cardiomyocytes (see Das *et al*., submitted elsewhere). Current amplitude was not different between genotypes (**Fig 1A-B**). As previously observed in an independent constitutive cardiac-specific knockout model (27), an inducible knockout model (25), a *Fgf13* hypomorphic line (26), and after *Fgf13* knockdown (19, 28), elimination of FGF13 hyperpolarized the V_1/2_ of VGSC SSI recorded from acutely isolated cardiomyocytes (V_1/2_ for WT, −82.4 ± 0.8 mV vs. −90.4 ± 0.9 mV for c*Fgf13^KO^*; **Fig 1C**). Also as previously reported (27), FGF13 ablation accelerated the rate of VGSC fast inactivation (Tau, **Fig 1D-E**). Since SSI depends primarily on the actions of membrane delimited voltage sensors, we investigated how FHFs restricted to the cytoplasm affect this property. Based on data in Das *et al*. showing that FGF13 regulates processes beyond Na_V_1.5 gating in cardiomyocytes and that FGF13 is in stoichiometric excess of Na_V_1.5 protein in heart lysates, we considered the possibility FGF13 exerted effects on VGSCs independent of FGF13’s interaction with Na_V_1.5.

**Figure 1:**
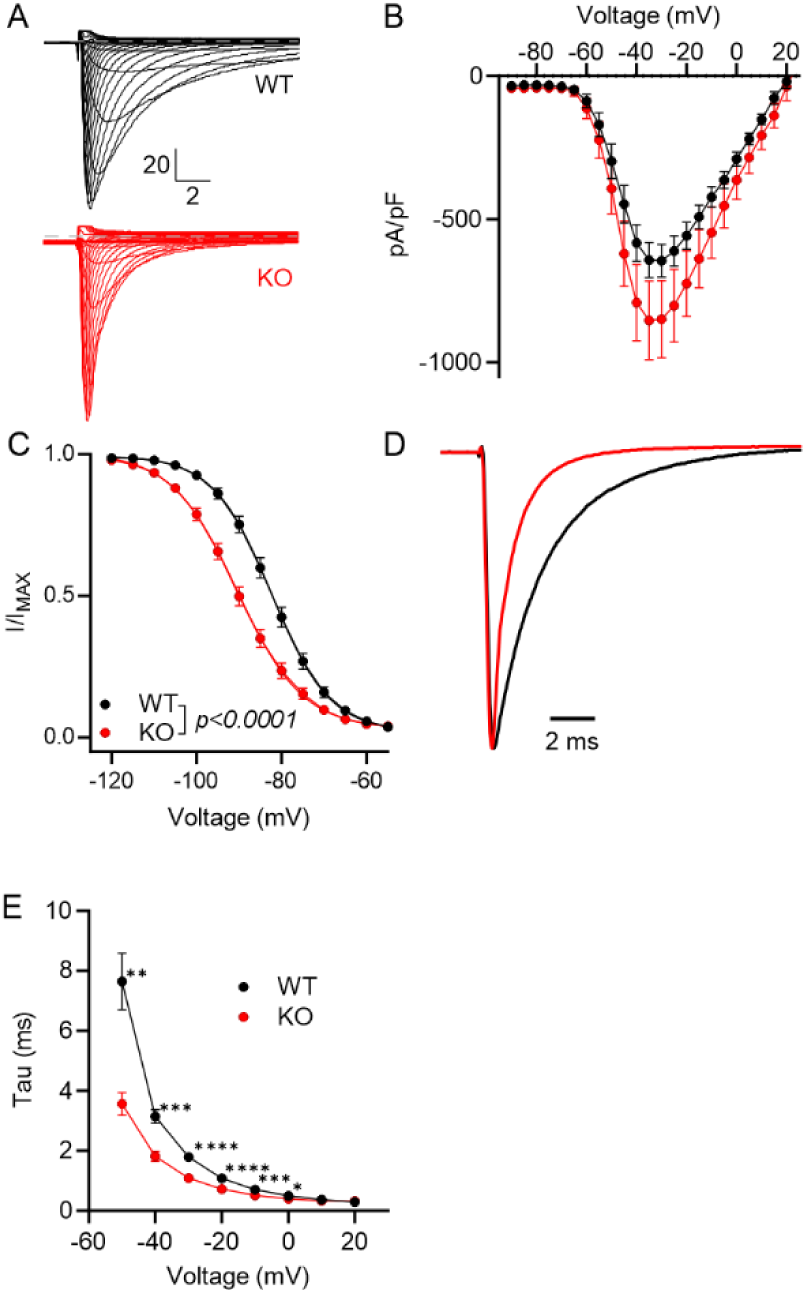
FGF13 regulates VGSC kinetics in cardiomyocytes. (A) Representative raw traces of sodium channel currents recorded from acutely isolated cardiac myocytes in response to depolarizing voltage steps from a holding potential of −120 mV. (B) Current-voltage relationship displaying peak current densities at various depolarizing voltages. (C) Steady-state inactivation curves showing normalized sodium currents elicited at −20 mV after conditioning from different holding potentials. (D) Representative raw traces of sodium currents at −20 mV, highlighting differences in fast inactivation kinetics. (E) Analyzed time constant of inactivation (Tau) for sodium currents recorded at different voltages. Statistical analysis: Two-way ANOVA with Bonferroni post-hoc tests (*p < 0.05, ***p<0.001, ****p<0.0001).

To test our hypothesis, we exploited previous structural studies (30, 31) and developed a strategy to replace FGF13 in cardiomyocytes with a mutant version incapable of binding to Na_V_1.5. The binding incompetent FGF13 bears an alanine substitution at Arg^120^ (“**FGF13^R/A^**”, **Fig 2A**), thus eliminating a side chain that inserts into a deep hole on the Na_V_1.5 C-terminal domain (**CTD**) surface with the channel’s His^1849^ at the hole’s base. Both residues are conserved among FHFs and VGSCs, respectively (**Sup Fig 1**), are critical for FHF-VGSC interaction (30): variants in either the FHF Arg or the Na_V_1.5 His residues disrupt binding and are associated with disorders characterized by abnormal VGSC function (32, 38). For these FGF13 expressing viruses, and in additional heterologous expression system studies, we use exclusively the Fgf13 “VY” splice variant, which is the dominant *Fgf13* transcript expressed in ventricle. Transcripts for the major neuronal splice variant, *Fgf13-S*, are <<10% of *Fgf13-VY* and FGF13-S protein is not detected in western blots of mouse ventricle (19). We confirmed that FGF13^R/A^ eliminated interaction with Na_V_1.5 in HEK293 cells by expressing FGF13 or FGF13^R/A^ with Na_V_1.5 and preforming immunoprecipitation of FGF13. **Fig 2B** shows that Na_V_1.5 co-immunoprecipitated with FGF13, but not FGF13^R/A^, analogous to previous data (39) obtained with a homologous FGF14 mutant and Na_V_1.6. With these results we were able to separate FGF13 regulation of VGSCs by direct interaction and by indirect means. We therefore virally expressed FGF13 or FGF13^R/A^ in ventricular cardiomyocytes isolated from c*Fgf13^KO^* for electrophysiological analyses and compared results to those also obtained from cardiomyocytes isolated from wild type mice or c*Fgf13^KO^* mice (**Fig 2C**). To allow for viral expression, recordings were performed 24-72 h after isolation and immediate infection. In c*Fgf13^KO^* mice, we confirmed efficient knockout of FGF13 and similar levels of expressed FGF13 or FGF13^R/A^ protein from their respective viruses (**Fig 2D**). VGSC current density was larger in c*Fgf13^KO^* cardiomyocytes infected with the FGF13^R/A^ virus but current densities in the other groups were indistinguishable from each other (**Fig 2E-F**).

**Figure 2:**
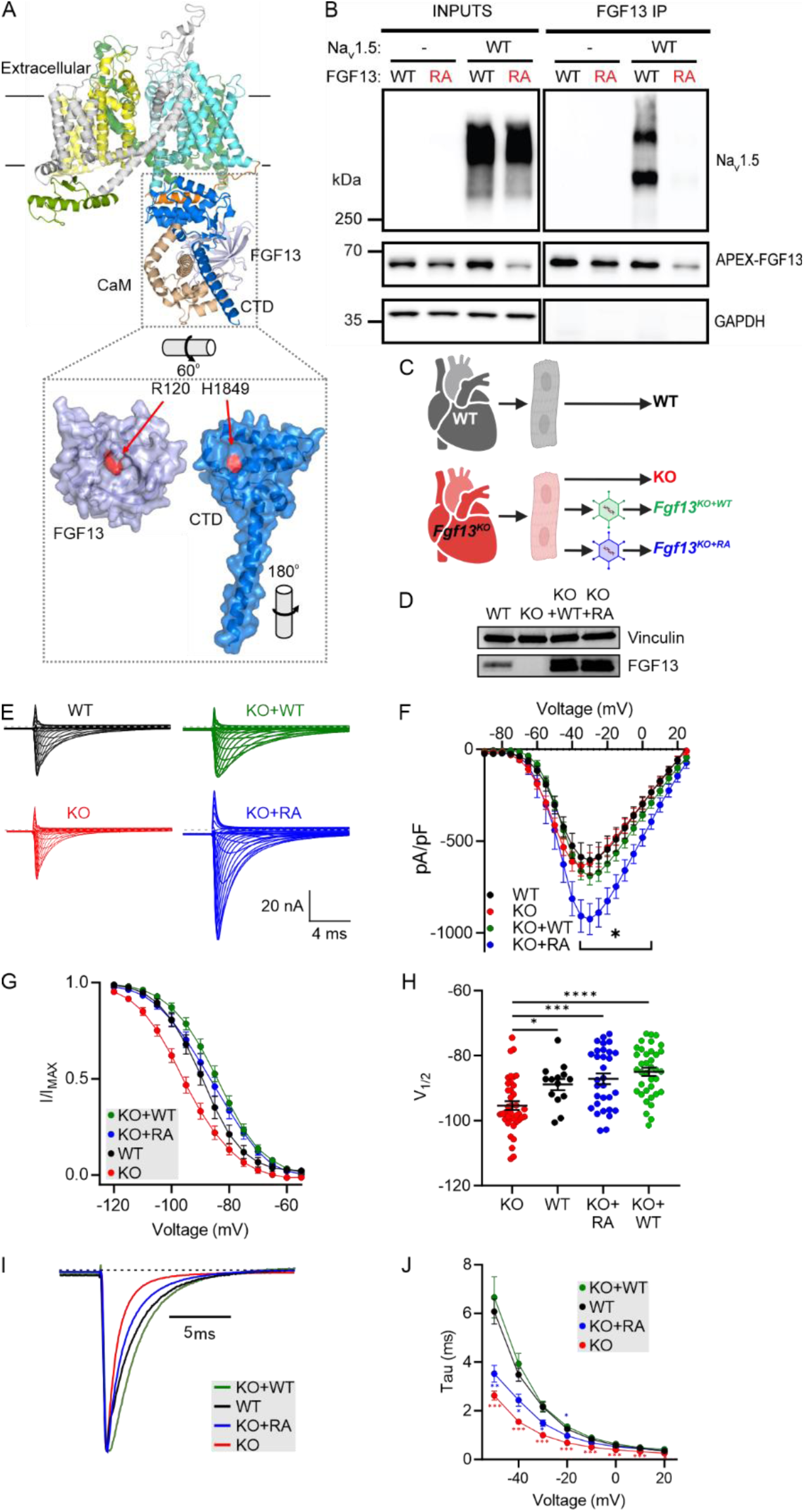
A binding incompetent FGF13 mutant confers a subset of WT-like regulatory functions in cardiomyocytes. (A) Overlay of the insect sodium channel crystal structure (PDB: 5X0M) with the Nav1.5 C-terminal domain (CTD) in complex with FGF13 (PDB: 4DCK), illustrating the FGF13 binding site on Nav1.5 CTD. (B) Immunoprecipitation of FGF13 in HEK cells co-expressing Nav1.5 and either wild-type (WT) or mutant (R120A) FGF13 (FGF13^R/A^), showing a loss of binding in FGF13^R/A^. (C) Experimental strategy to re-express FGF13 or FGF13^R/A^ in cardiac myocytes from FGF13 knockout (KO) mice using AAV vectors. (D) Western blot confirming the expression of FGF13 and FGF13^R/A^ in KO myocytes post-rescue. (E) Representative sodium current traces from all experimental groups. (F) Current-voltage relationships showing peak current densities. (G) Steady-state inactivation curves displaying normalized currents at −20 mV. (H) Half-maximal voltage (V_1/2_) of inactivation showing rescue effects of FGF13 and FGF13^R/A^. (I) Representative sodium current traces at −20 mV illustrating differences in fast inactivation. (J) Time constants (Tau) of inactivation at varying voltages. Statistical analysis: Two-way ANOVA with Bonferroni post-hoc tests (*p < 0.05, **p < 0.01, ***p < 0.001).

We then queried SSI and kinetics of fast inactivation. As in acutely isolated cardiomyocytes, the V_1/2_ for SSI in WT was significantly depolarized compared to the V_1/2_ for SSI in c*Fgf13^KO^* cardiomyocytes (**Fig 2G-H**). Expression of FGF13 in c*Fgf13^KO^* cardiomyocytes significantly depolarized the V_1/2_ of SSI, demonstrating successful rescue. Similarly, re-expression of the binding incompetent FGF13^R/A^ depolarized the V_1/2_ of SSI (**Fig 2G-H**), suggesting that FGF13 binding to Na_V_1.5 is not necessary for regulation of SSI. Also as in acutely isolated cardiomyocytes, VGSC currents in cultured c*Fgf13^KO^* cardiomyocytes showed faster inactivation than in WT cardiomyocytes. Viral expression of FGF13 in c*Fgf13^KO^*cardiomyocytes slowed VGSC inactivation, but viral expression of FGF13^R/A^ did not (**Fig 2I-J**). Thus, fast inactivation requires interaction between FGF13 and the channel’s C-terminal domain. We obtained concordant results with a heterologous expression system. In HEK293 cells, we expressed Na_V_1.5 with FGF13 (either WT or FGF13^R/A^) and GFP; or with GFP only. Expression of Na_V_1.5 with GFP only is analogous to *Fgf13^KO^* in cardiomyocytes since HEK293 cells do not express endogenous FGF13, while expression of Na_V_1.5 with FGF13 corresponds to WT cardiomyocytes. As in cardiomyocytes, Na_V_1.5 SSI was hyperpolarized, and Tau was faster in the absence of FGF13 (GFP expression only) than with FGF13. Also as in cardiomyocytes, expression of the binding incompetent FGF13^R/A^ restored SSI but not Tau (**Sup Fig 2A-E**).

How does FGF13 affect Na_V_1.5 SSI via a binding-independent mechanism? Since SSI is generally regulated by the membrane delimited voltage sensors and that—having excluded a direct interaction mechanism—a hyperpolarizing shift in VGSC SSI is near-pathognomonic for a reduction in membrane stiffness via depletion of accessible membrane cholesterol (40–42), we considered biochemical data showing that FGF13 regulates local membrane cholesterol content through interaction with cavins (43), a set of caveolae regulator proteins (44), without affecting the total cholesterol pool. Specifically, we hypothesized that FGF13 affected local membrane cholesterol, independent of binding to Na_V_1.5, to regulate SSI of Na_V_1.5 in cardiomyocytes. As cholesterol regulation of VGSC gating may be channel specific as suggested by differing effects after cholesterol depletion (40–42, 45), we first tested whether manipulation of membrane cholesterol affected Na_V_1.5 channels. Indeed, depletion of membrane cholesterol with MβCD induced a hyperpolarizing shift in the V_1/2_ of SSI for Na_V_1.5 (co-expressed with GFP as a control) and for Na_V_1.5 co-expressed with FGF13, albeit to a lesser extent (**Sup Fig 3A-B**). In contrast, addition of cholesterol depolarized SSI for Na_V_1.5 (co-expressed with GFP) but did not affect SSI for Na_V_1.5 co-expressed with FGF13 (**Sup Fig 3C-D**).

Further, we showed that FGF13 affected membrane cholesterol by exploiting ALOD4, a recombinant domain of the soluble bacterial toxin anthrolysin O (ALO) that binds accessible plasma membrane cholesterol (46, 47) to visualize effects in HEK293 cells. ALOD4 labels the small pool of membrane cholesterol (<10% of total) that is metabolically active (48) and most accessible to cholesterol depleting reagents such as cyclodextrins. Application of ALOD4 to live cells labeled the plasma membrane accessible cholesterol pool, as observed by fluorescent imaging of HEK293 cells (**Fig 3A** and **Sup Fig 4A**). Incubation with MβCD (500 µM for 60 min) depleted the ALOD4 fluorescent signal while addition of cholesterol augmented the signal (**Fig 3A** and **Sup Fig 4A**), showing specificity of ALOD4. We then queried whether the presence of FGF13 or FGF13^R/A^ (compared to GFP control) affected the ALOD4 signal. While in control GFP-transfected cells the ALOD4 signal in maximum projection confocal stacks appeared evenly distributed to the cell periphery, with FGF13 or FGF13^R/A^ transfection we observed irregular signal scattered throughout the image, consistent with the presence of membrane ruffles (**Fig 3B** and **Sup Fig 4B-C**), which are regulated by local changes in cholesterol (49). Despite the difference in the pattern of fluorescent signal, the integrated total signal was not different among the three groups (**Fig 3C**), reflecting expected cellular homeostatic mechanisms (50) and consistent with previous data showing that total cholesterol in heart lysates was unperturbed in *Fgf13* knockout mice despite a change in local membrane cholesterol distribution (43). Consistent with the microscopy images, we prepared cellular lysates and quantified ALOD4 protein by gel electrophoresis and in-gel imaging of the ALOD4 fluorescence signal. **Fig 3D-E** shows ALOD4 signal depletion by MβCD within all groups and demonstrates that the overall accessible cholesterol is unaffected by expression of FGF13 or FGF13^R/A^. Together, these data show that FGF13 does not affect total cell cholesterol, but does alter the distribution (compared to no FGF13) of accessible cholesterol in the membrane, consistent with previous biochemical data (43). Moreover, as HEK293 cells do not express endogenous Na_V_1.5, these data (along with data showing similar effects with the Na_V_1.5 binding incompetent FGF13^R/A^) suggest that the effects of FGF13 on membrane cholesterol are independent of Na_V_1.5 binding.

**Figure 3:**
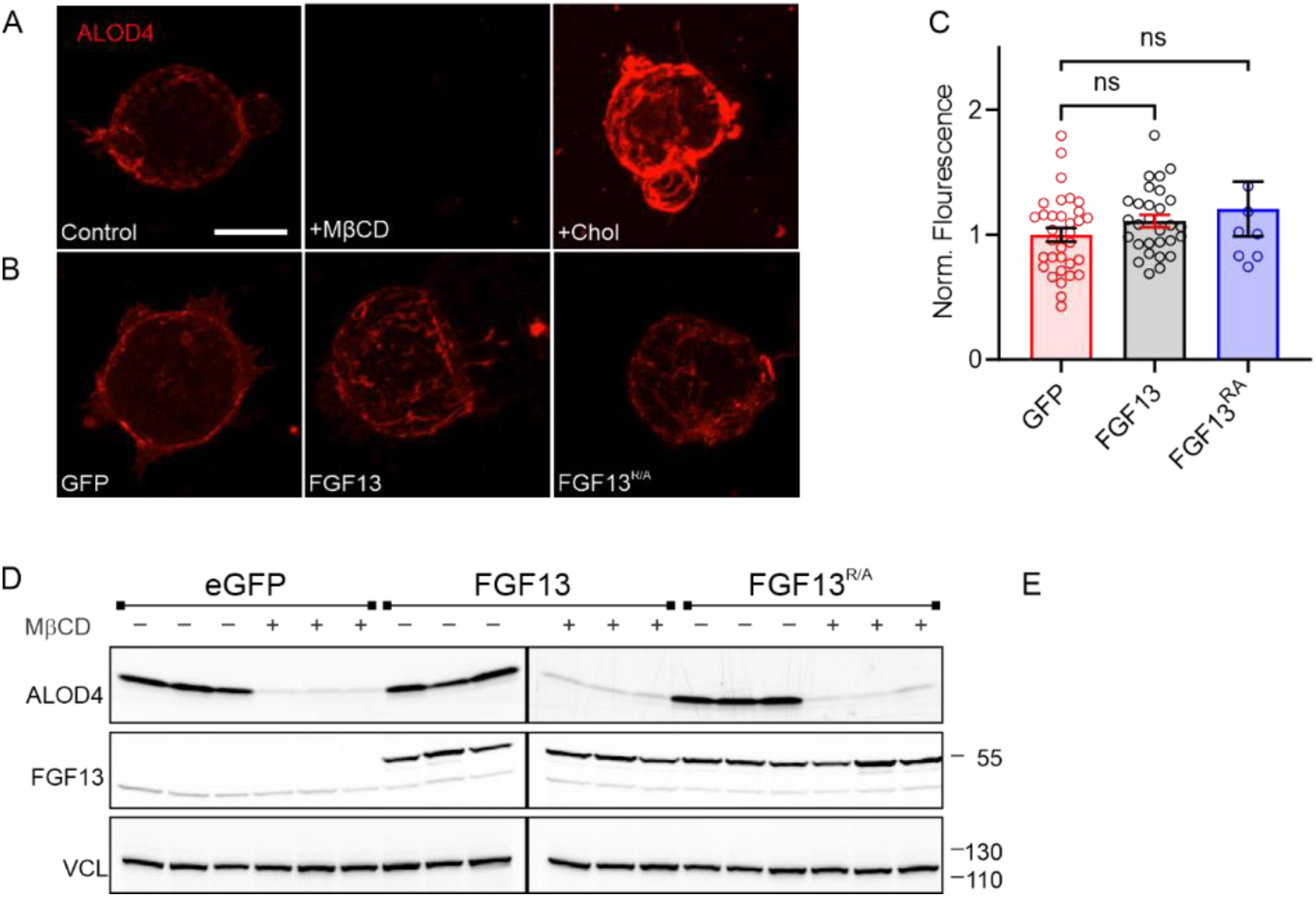
ALOD4 detects membrane accessible cholesterol. Confocal images of ALOD4 staining in HEK cells under control, MβCD-treated, and cholesterol-enriched conditions (Scalebar – 10 µm) (A) and after expression of GFP, FGF13, and FGF13^R/A^ (B). (C) Quantification of ALOD4 intensity, showing no significant differences between FGF13 and FGF13^R/A^-expressing cells. Statistical analysis: One-way ANOVA. (D, E) In-gel fluorescence imaging of ALOD4 showing no difference total membrane accessible cholesterol in HEK cells expressing FGF13 or FGF13^R/A^ or GFP.

We then applied ALOD4 to isolated cardiomyocytes. In WT cells, accessible cholesterol is polarized, with concentration at the IDs, as measured by line scans (starting 1 µm before the cell edge) perpendicular to the cardiomyocyte short axis (**Fig 4A-E**, **Sup Fig 5A**). In contrast, in cardiomyocytes from c*Fgf13^KO^* mice the signal intensity from the ID was reduced (signal between 2-5 µm along the line) and relatively more signal was observed throughout the rest of the cell (measured between 6-20 µm along the line). We demonstrated the specificity of the ALOD4 signal in cardiomyocytes by showing that cholesterol depletion with MβCD reduced the signal in WT cardiomyocytes while addition of cholesterol increased the signal (**Sup Fig 5B**). Despite the genotype differences in pattern and concentration of signal at the ID (**Fig 4D**), along 20 µm line scans there were no differences in area under the curve measurements (**Fig 4C**), suggesting that the total cholesterol was not different between genotypes. Indeed, quantifying total cholesterol with filipin fluorescence showed no difference (**Fig 4F-G**). We confirmed the specificity of the filipin signal by showing its depletion after application of MβCD (**Fig 4F-G**, **Sup Fig 5C**). Together, these data showing that ablation of FGF13 drives redistribution of accessible membrane cholesterol from the cardiomyocyte ID to the midsection without affecting the total cholesterol pool are consistent with previous biochemical data showing that FGF13 ablation redistributes membrane cholesterol (43).

**Figure 4:**
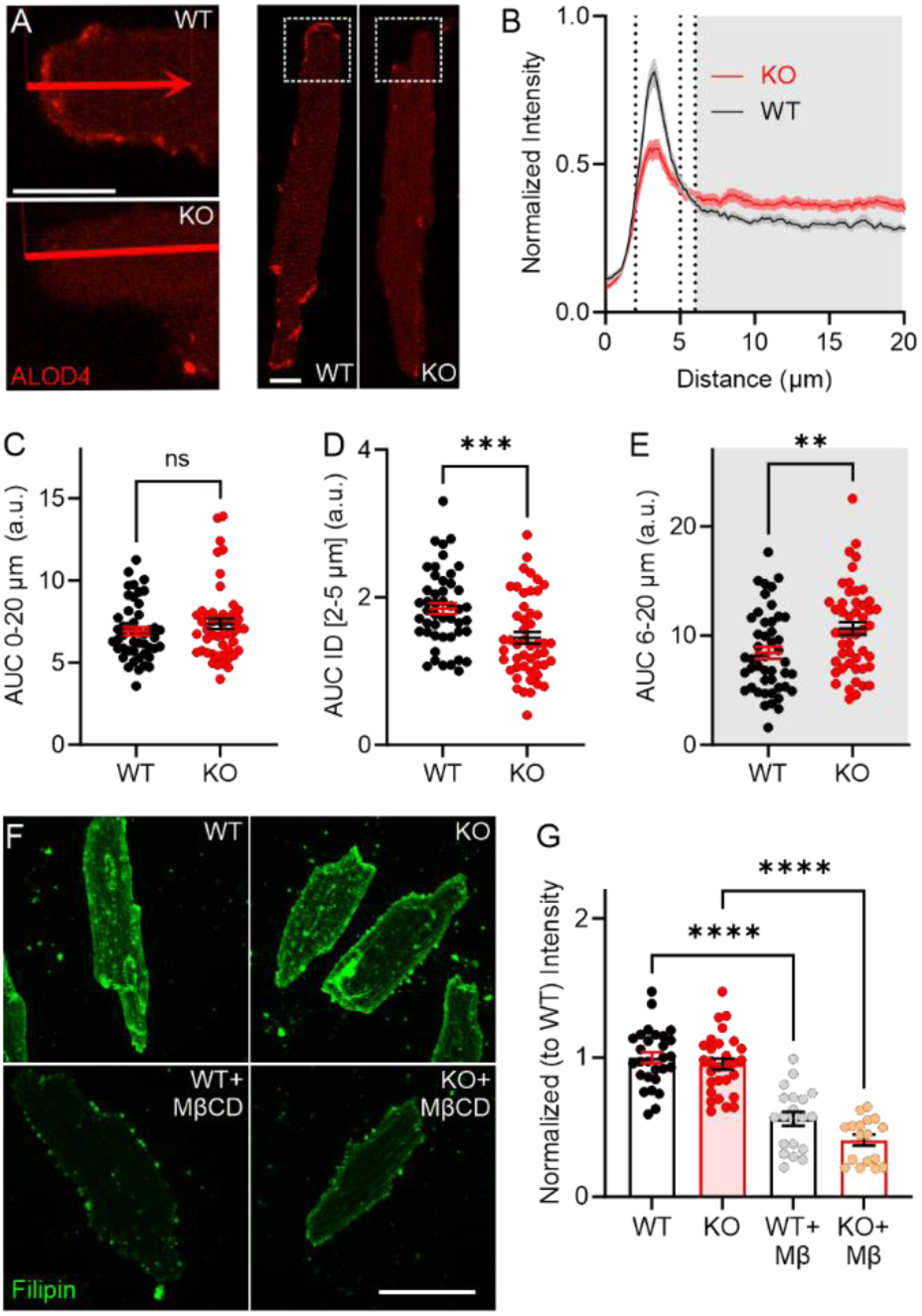
Accessible membrane cholesterol is concentrated at the intercalated discs in cardiomyocytes and is regulated by FGF13. (A) Confocal images of ALOD4 staining in c*Fgf13^KO^* cardiac myocytes, showing altered distribution compared to WT (Scalebars – 10 µm). (B) Quantification of ALOD4 signal from intercalated discs to the lateral membrane, highlighting localization differences in c*Fgf13^KO^* myocytes. (C-E) Quantification of ALOD4 intensity distribution, showing no significant change in total signal intensity. Statistical analysis: Unpaired t-test (**p<0.01, ***p<0.001). (F) Filipin staining of cardiac myocytes, demonstrating overall cholesterol depletion following MβCD treatment (Scalebar – 50 µm). (G) Quantification of filipin signal intensity in WT and KO myocytes, showing no cholesterol differences. Statistical analysis: One-way ANOVA with Bonferroni post-hoc tests (**p < 0.01).

Not only did *Fgf13* knockout affect membrane cholesterol distribution, but it also affected the amount of Na_V_1.5 at the ID. **Fig 5A** shows immunocytochemistry for Na_V_1.5 in cardiomyocytes isolated from WT or c*Fgf13^KO^* mice. Co-staining with an antibody for N-cadherin (N-Cad) marks the intercalated disc, where one of the two major Na_V_1.5 channel pools resides. Fluorescence intensity line scans perpendicular to the cardiomyocyte short axis and normalized to the signal in WT cells show a peak Na_V_1.5 signal in WT at the ID (overlapping with the N-Cad signal, **Fig 5A-B**). The Na_V_1.5 fluorescence intensity in c*Fgf13^KO^* cells was 39.9 ± 14.6% lower (**Fig 5C**, quantified by area under curve measurements for the first two micrometers of the line scan, **Fig 5D**). We considered that the reduction in Na_V_1.5 at the ID in c*Fgf13^KO^* cells resulted from loss of FGF13 regulated trafficking of Na_V_1.5 to the ID, as previously hypothesized (43), and/or from reduction in local membrane cholesterol, as in **Fig 4**. To test if a reduction in local membrane cholesterol was sufficient to affect Na_V_1.5 at the IDs, we treated cardiomyocytes isolated from WT mice and observed that treatment with MβCD reduced the Na_V_1.5 fluorescence intensity at the ID by 18.6 ± 6.5% (**Fig 5E-G**). Together, these data indicate that FGF13 ablation not only reduces Na_V_1.5 at the ID but that perturbation of membrane accessible cholesterol by FGF13 ablation is a likely mechanism. Based on these imaging data, we hypothesized that known ID components would be depleted from Na_V_1.5 immunoprecipitates in c*Fgf13^KO^* hearts compared to WT hearts. We therefore identified by mass spectrometry proteins co-immunoprecipitated with Na_V_1.5 from WT or c*Fgf13^KO^* hearts (n=3 each). Principal component analysis and Pearson correlation statistics showed that the set of proteins co-immunoprecipitated from WT hearts was distinct from the set co-immunoprecipitated from c*Fgf13^KO^*hearts (**Sup Fig 6A-B**). We detected 74 co-immunoprecipitated proteins depleted (*P*<0.05; log_2_FC≥1.0) in KO hearts compared to WT hearts (**Fig 6A** and **Supp Table 1**) and, indeed, these included several ID components (e.g., proteins encoded by *Gja1*, *Ctnnb1*, *Cdh2*, *Dlg1*, and *Vcl*). Further, gene set enrichment analysis (**Fig 6B-D**) showed that top terms included “fascia adherens” (GO:0005916) and “intercalated disc” (GO:0014704), terms not found in the analysis of the set of co-immunoprecipitated proteins enriched in c*Fgf13^KO^* hearts (**Supp Table 1**). Also notable was the identification of calmodulin (*Calm1*) as one the co-immunoprecipitated proteins depleted in KO hearts compared to WT hearts (log_2_FC=1.63, *P*=0.02) since a previous study reported that FGF13 increases the affinity between Na_V_1.5 and calmodulin (51).

**Figure 5:**
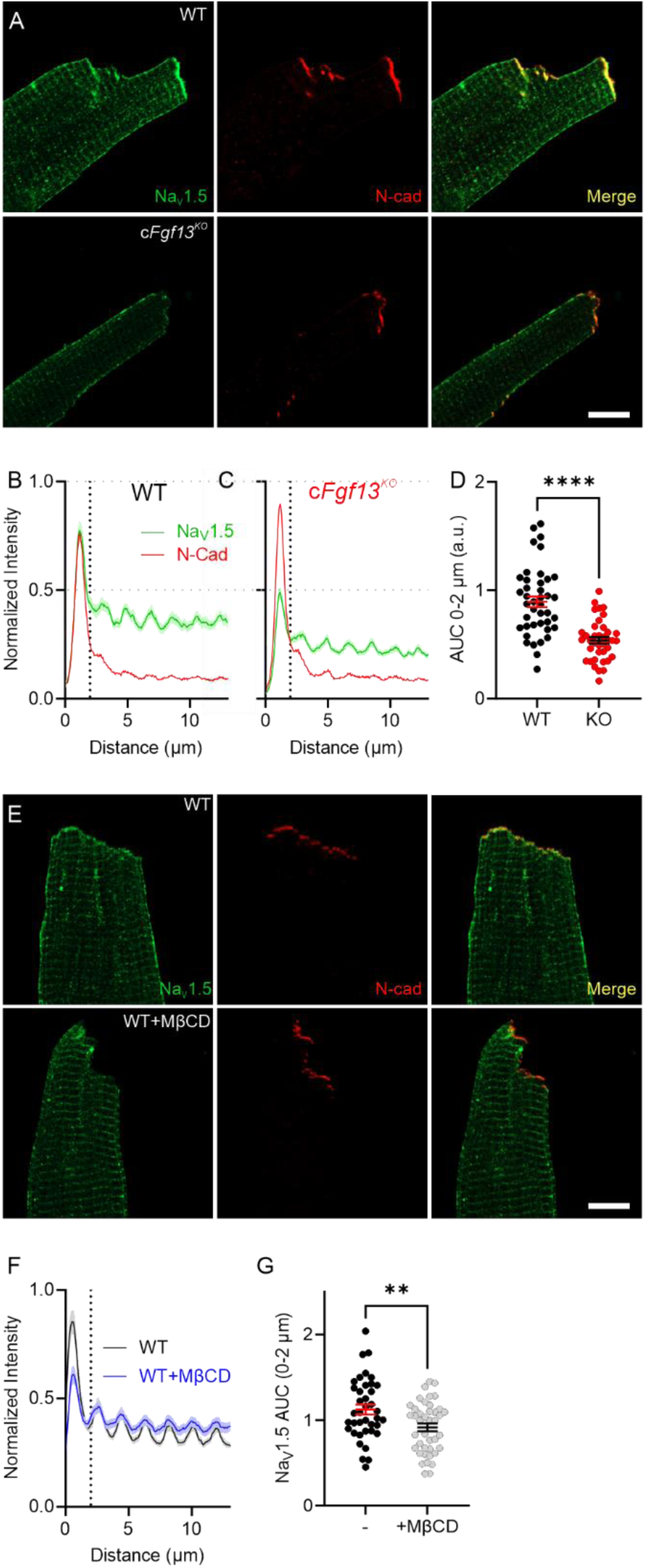
FGF13 and cholesterol regulate Na_V_1.5 at the intercalated disc. (A) Confocal images showing Nav1.5 localization in cardiac myocytes from WT and c*Fgf13^KO^* KO mice (Scalebar – 10 µm). (B, C) Quantification of Nav1.5 signal intensity at the intercalated disc and lateral membrane in WT and KO myocytes. (D) Total Nav1.5 signal intensity in KO myocytes, showing a significant reduction. Statistical analysis: Unpaired t-test (****p<0.0001) (E) Confocal images of Nav1.5 in WT myocytes under control and MβCD-treated conditions (Scalebar – 10 µm). (F, G) Quantified Nav1.5 intensity at the intercalated disc and total levels after MβCD treatment, resembling the c*Fgf13^KO^* phenotype. Statistical analysis: Unpaired t-test (**p < 0.01).

**Figure 6:**
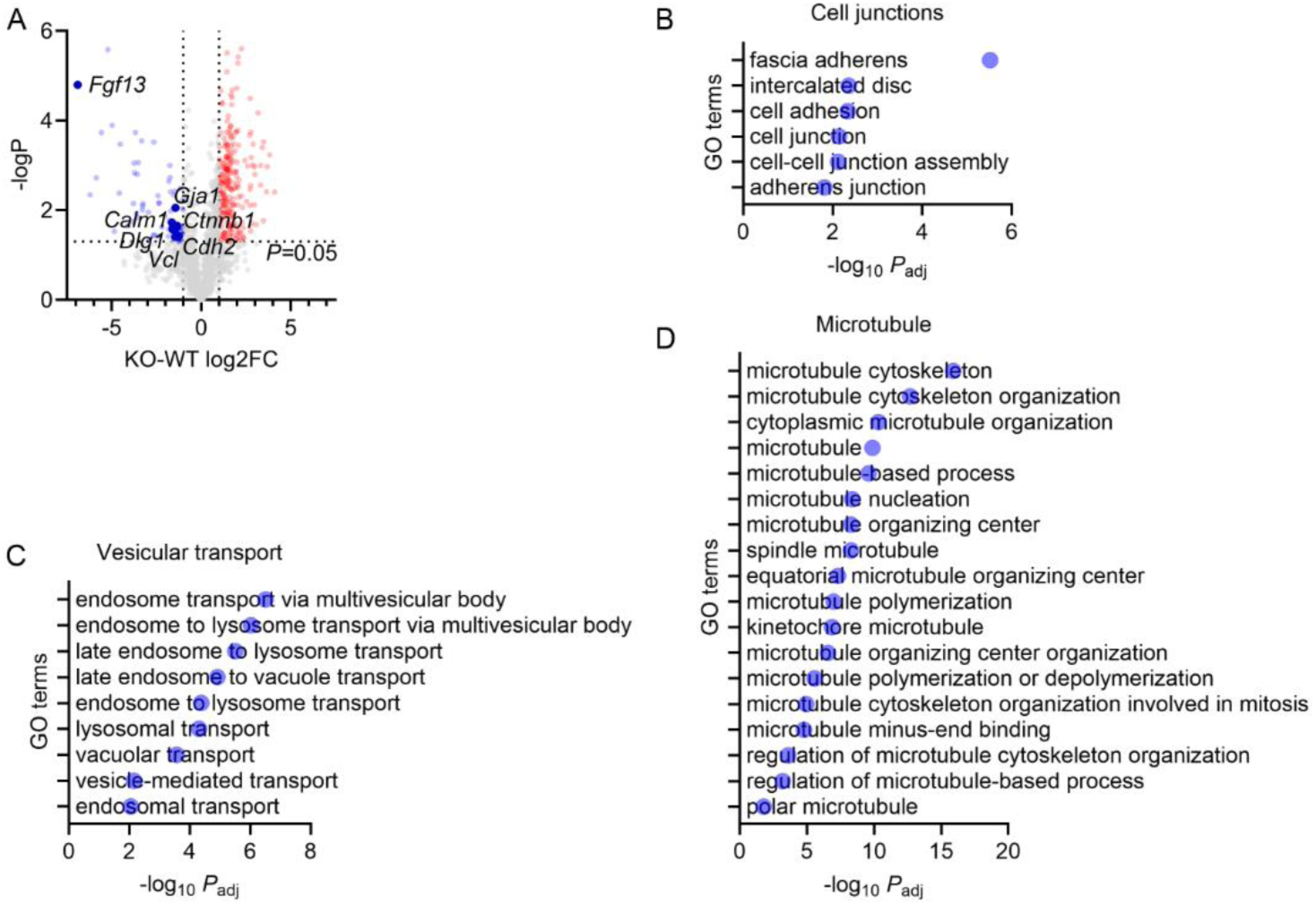
Knockout of *Fgf13* affects the localization of Na_V_1.5 at the intercalated disc. (A) Volcano plot of proteins co-immunoprecipitated with Na_V_1.5 from WT or c*Fgf13^KO^* hearts. Key proteins are highlighted. (B-D) Gene set enrichment analyses of proteins enriched in co-immunoprecipitates of Na_V_1.5 in WT (compared to c*Fgf13^KO^* hearts).

Moreover, consistent with our accompanying manuscript showing that FGF13 regulates Cx43 trafficking in vesicles along microtubules (Das *et al*.), two other sets of top gene ontology terms found in the analysis of co-immunoprecipitated proteins depleted from WT hearts were terms associated with vesicular transport and microtubules (**Fig 6C-D** and **Supp Table 1**). None of these terms were found in the analysis of the set of proteins enriched in the co-immunoprecipitates from c*Fgf13^KO^* hearts. Thus, these data underscore that FGF13 affects the subcellular distribution of Na_V_1.5 and provide mechanistic insight into the altered Na_V_1.5 subcellular localization observed in c*Fgf13^KO^* hearts.

The redistribution of Na_V_1.5 away from the ID in the absence of FGF13 could affect VGSC properties based on a previous study showing differences in VGSC amplitude and SSI depending on where on the cardiomyocytes recordings were performed (52). Currents recorded from the ID were larger and the V_1/2_ of SSI was depolarized compared to currents recorded from the midsection (lateral membrane, LM). We hypothesized that the change in subcellular distribution of Na_V_1.5 as regulated by FGF13 was an important contributor. We therefore performed macropatch recordings from the ID and midsection, which differed in WT vs. c*Fgf13^KO^* cardiomyocytes. In WT cardiomyocytes, peak currents were larger at the ID than at the midsection, as expected, yet that differential was lost in c*Fgf13^KO^* cells and the current amplitudes at either the ID or midsection were not different from the midsection in WT cells (**Fig 7A-C**). The macropatch data also suggest that FGF13 contributes to the previously observed depolarization of the V_1/2_ of SSI at the ID compared to the midsection. In WT cells, SSI at the ID was depolarized compared to V_1/2_ at the midsection, but in c*Fgf13^KO^* cardiomyocytes there was no statistically significant difference (**Fig 7 D-F**). Using sensitivity to tetrodotoxin (TTX), the previous study suggested that the amplitude and SSI differences between the ID and midsection reflected predominantly TTX-resistant Na_V_1.5 channels expressed at the ID and TTX-sensitive neuronal VGSCS at the midsection (52). Here, we find that SSI of VGSCs at the ID and the midsection in WT cells were depolarized compared to their counterparts in c*Fgf13^KO^* cardiomyocytes. Thus, FGF13 affects SSI of VGSCs independent of channel location, suggesting that FGF13 affects both Na_V_1.5 and neuronal VGSCs expressed in cardiomyocytes, consistent with conserved binding determinants among all VGSCs (**Sup Fig 2**).

**Figure 7:**
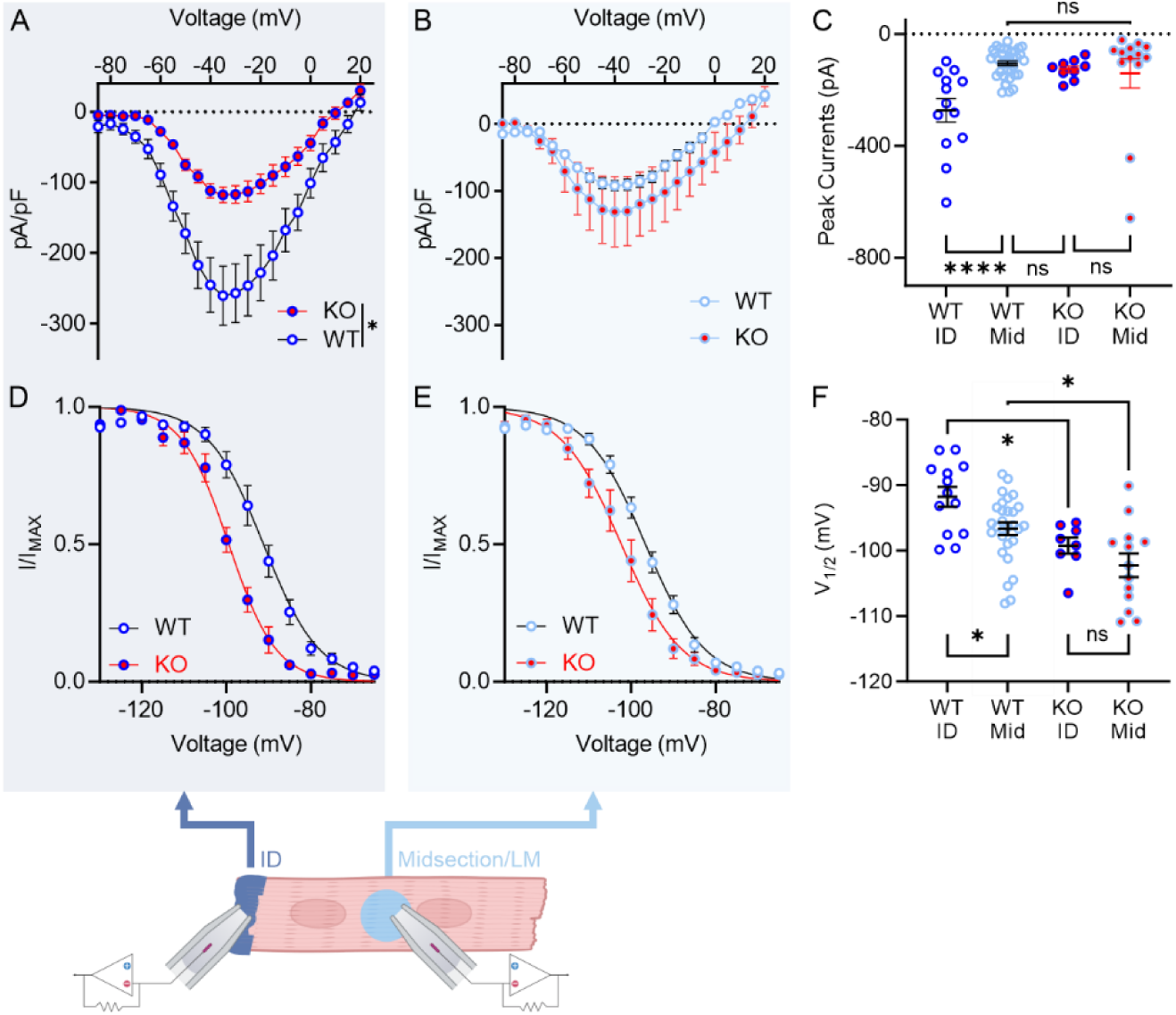
Macropatch experiments show loss of VGSC currents at the intercalated disc of cardiomyocytes from c*Fgf13^KO^* hearts. Current voltage relation curves from intercalated discs (ID) (A) and lateral membranes (LM) (B) of WT and KO myocytes. Steady-state inactivation curves from ID (D) and LM (E) of WT and KO myocytes. (E) Experimental design for macropatch clamp experiments. (F) Peak sodium current densities showing differences between ID and LM of WT but not KO myocytes. One-way ANOVA with Bonferroni post-hoc tests (****p < 0.0001). V_1/2_ of inactivation values, showing significant differences between WT and KO myocytes at ID (F) and LM (G). Statistical analysis: Unpaired t-test (***p < 0.001). V_1/2_ of inactivation values, showing significant differences between ID and LM of WT (H) but not KO (I) myocytes. Statistical analysis: Unpaired t-test (*p < 0.05).

Since *Fgf13* ablation affects the polarized distribution of membrane accessible cholesterol, which regulates SSI in HEK293 cells (**Sup Fig 3**), we hypothesized that the lack of difference between the V_1/2_ of SSI at the ID and midsection in c*Fgf13^KO^* cardiomyocytes (compared to WT cells) was due to the resulting perturbation of membrane cholesterol. Technical limitations prevented measurements of SSI in macropatch experiments after depleting or adding membrane cholesterol, but since we observed that FGF13 affected SSI at either the ID or midsection, we recorded SSI in cardiomyocytes in the whole cell configuration, which was amenable to cholesterol manipulation. Consistent with results in HEK293 cells, depletion of cholesterol with MβCD hyperpolarized the V_1/2_ of SSI recorded in WT or c*Fgf13^KO^*cardiomyocytes (**Fig 8A-B**). Addition of cholesterol to c*Fgf13^KO^* cardiomyocytes depolarized the V_1/2_ of SSI but did not affect the V_1/2_ of SSI in WT cardiomyocytes (**Fig 8C-D**). Again, results in HEK293 cells were analogous and together suggest that the effects of cholesterol on SSI in WT cardiomyocytes are saturated.

**Figure 8:**
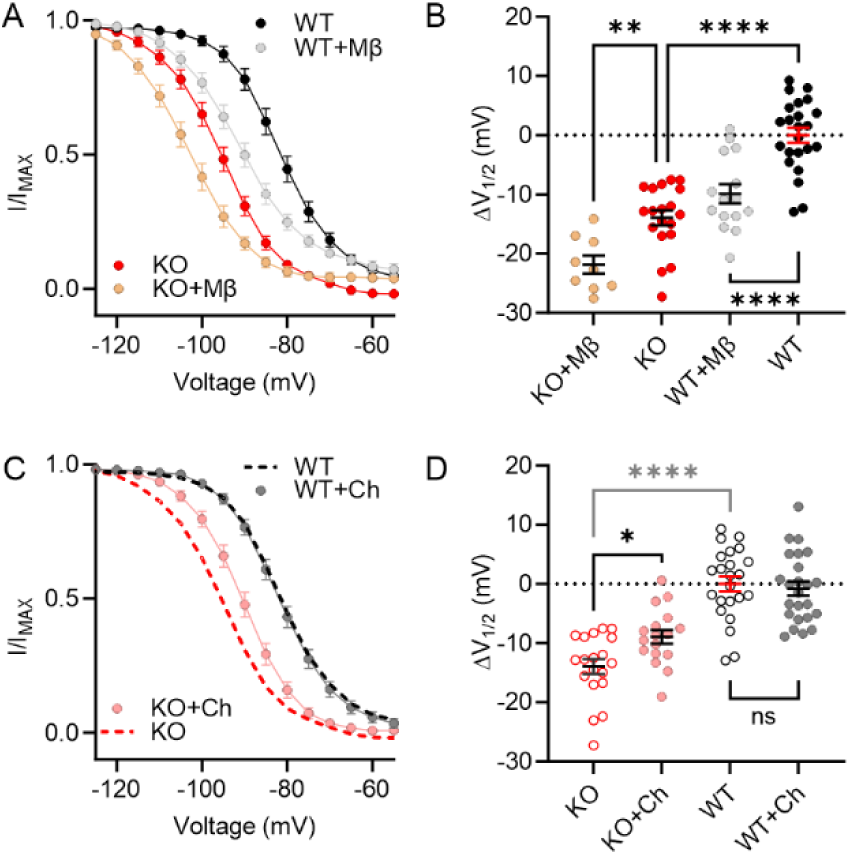
FGF13 differentially affects consequences of manipulation of membrane accessible cholesterol affects SSI of VGSCs in cardiomyocytes. Steady-state inactivation curves of sodium currents from WT and KO myocytes under MβCD treated (A) and cholesterol enriched (C) conditions. Changes in the V_1/2_ of inactivation values showing MβCD treatment (B) and cholesterol enrichment (D) induced effects. Data for WT and KO are repeated in panels (C) and (D). Statistical analysis: One-way ANOVA with Bonferroni post-hoc tests (*p < 0.05, **p < 0.01, ****p < 0.0001).

Beyond affecting SSI, a previous report showed that *Fgf13* ablation in cardiomyocytes reduced VGSC currents at elevated temperatures, leading to a temperature dependent conduction block (26); analogous data from neuron-specific *Fgf13* knockout models showed reduced heat nociception in dorsal root ganglion neurons (mediated by Na_V_1.7 channels) (53, 54). Since cholesterol affects membrane fluidity in a temperature dependent fashion, we hypothesized that the observed conduction block was driven at least in part by the FGF13 dependent change in membrane cholesterol. To test our hypothesis, we first asked if the effects of temperature on Na_V_1.5 channels was independent of a direct interaction between FGF13 and Na_V_1.5 by performing paired recordings at 25 °C and 40 °C in HEK293 cells expressing Na_V_1.5 and FGF13 (or GFP as a control). We observed a marked reduction of Na_V_1.5 current amplitude at 40 °C (**Sup Fig 7A-B**). In contrast, current amplitude at 40 °C was not reduced when FGF13 or the binding incompetent FGF13^R/A^ were co-expressed (**Sup Fig 7C-F**), showing that FGF13 preserves VGSC current amplitude at elevated temperatures independent of FGF13 binding to the channel. Similar to the protection afforded by co-expression of either FGF13 or FGF13^R/A^, we observed that addition of cholesterol to the membrane was also protective (**Sup Fig 7G-H**). The addition of FGF13, FGF13^R/A^, or cholesterol all increased the Q_10_ above that observed in cells expressing Na_V_1.5 and GFP only. We obtained congruent data in ventricular cardiomyocytes, observing no reduction in current amplitude when the bath temperature for WT cells increased from 25 °C to 40 °C (**Fig 9A-B**), but a significant reduction of current amplitude of c*Fgf13^KO^*cells in which the bath temperature was raised to 37 or 40 °C (**Fig 9C-D**). Moreover, viral expression of either WT FGF13 or the binding incompetent FGF13^R/A^ in cardiomyocytes isolated form c*Fgf13^KO^* animals prevented to the reduction in current amplitude at elevated temperatures (**Fig 9E-H**), demonstrating that the expression of FGF13 can acutely provide temperature stability to VGSC currents in cardiomyocytes and that the effect of FGF13 is independent of Na_V_1.5 binding. Since cholesterol and temperature both affect membrane fluidity, we tested if addition of cholesterol to c*Fgf13^KO^* cardiomyocytes stabilized VGSC currents as temperature increased. We added cholesterol to c*Fgf13^KO^*cells and then recorded VGSC current amplitudes while raising the bath temperature. Similar to viral expression of FGF13, addition of cholesterol prevented the reduction in current amplitude observed in c*Fgf13^KO^*cardiomyocytes (**Fig 9I-J**). Thus, like FGF13 (present in WT cardiomyocytes) or after viral expression of FGF13 or FGF13^R/A^ in c*Fgf13^KO^* cardiomyocytes, cholesterol protects VGSC current at elevated temperatures (**Fig 9K**) and increased the Q_10_ compared to the values obtained in c*Fgf13^KO^* cardiomyocytes (**Fig 9L**).

**Figure 9:**
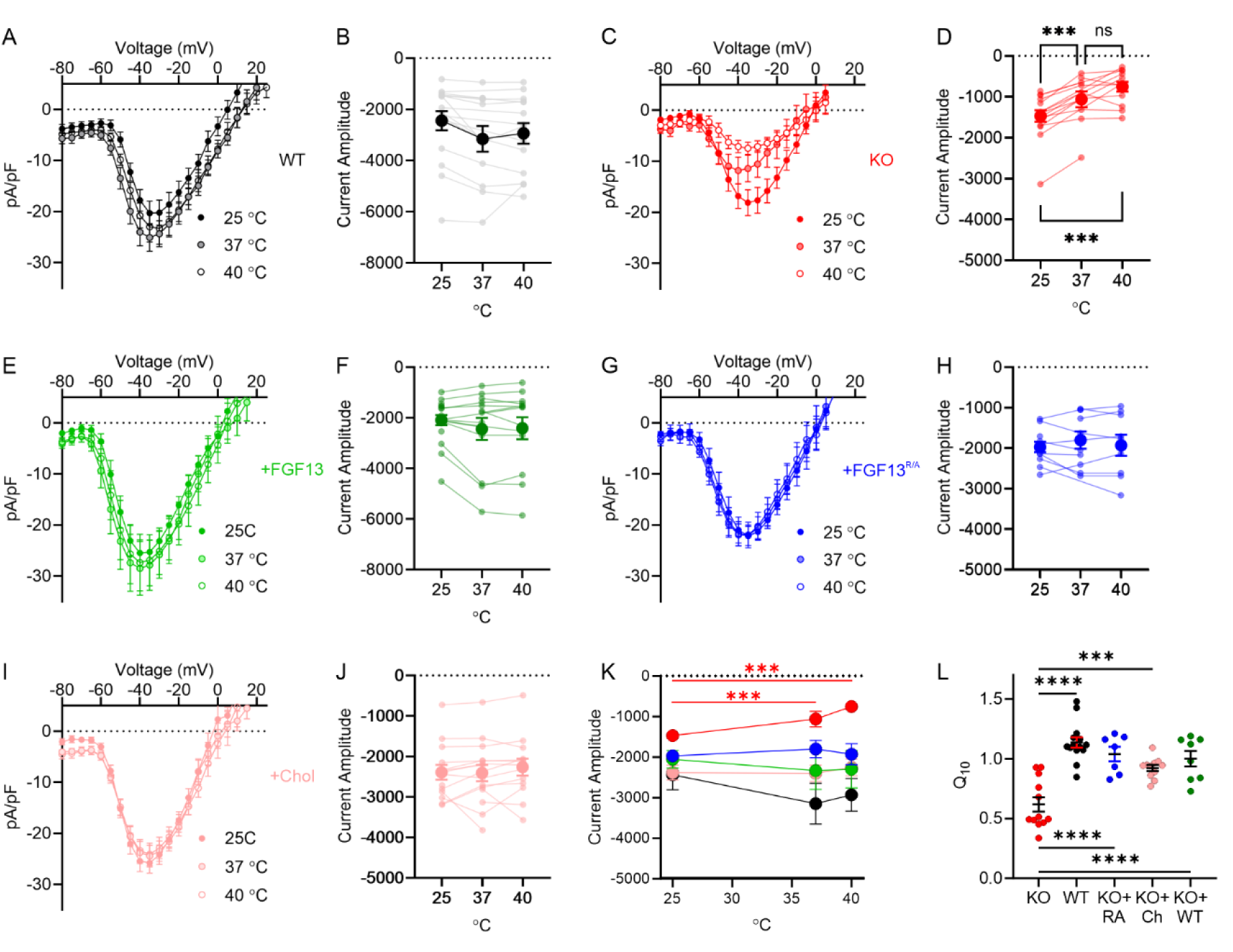
FGF13 and cholesterol protect VGSC currents at elevated temperatures in cardiomyocytes. Current-voltage relationships for sodium currents in WT (A), KO (C), KO + FGF13 (E), KO + FGF13^R/A^ (RA) (G), and KO + cholesterol enriched (I) groups. Peak current changes in response to increased temperature in WT (B), KO (D), KO + WT FGF13 (F), KO +FGF13^R/A^ (H), and KO + cholesterol enriched (J) groups. One-way ANOVA with Bonferroni post-hoc tests (***p < 0.001). (K) Summary data of peak current density changes with temperature. (L) Calculated Q_10_ for the tested conditions. Statistical analysis: One-way ANOVA with Bonferroni post-hoc tests (***p < 0.001).

## Discussion

Our data redefine and expand the roles for members of the FHF subfamily of FGFs and offer insights into how channel auxiliary proteins affect the function of their regulatory targets, the channel pore forming subunits. In cardiomyocytes, previous studies focused almost exclusively on how these FHFs serve as VGSC binding partners and regulate VGSC (8, 17, 19–22, 24–28, 32, 38, 43, 51, 55–58). This focused paradigm followed a yeast two hybrid screen that identified FGF12 as an interactor with the cytoplasmic C-terminus of Na_V_1.9 (*SCN11A*) and was bolstered by studies that discovered variants in a neuronal FHF associated with ataxias and variants in cardiomyocyte FHFs with arrhythmias. Observations of perturbed VGSC function when channels were co-expressed with disease associated variants provided pathogenic mechanisms to bolster that paradigm (20, 22–24, 59, 60). Crystal structures showing the direct interaction between FHFs and the cytoplasmic C-termini of VGSCs (30, 31, 61, 62) provided additional support. Building upon the identification of FHFs as regulators of Na_V_1.5 trafficking and targeting in cardiomyocytes (19, 25), roles in the trafficking and targeting of other ion channels have also been proposed after the observation that FHFs affect currents beyond those of VGSCs (25, 63).

As FHFs are members of the FGF superfamily, however, VGSC regulation is not *a priori* an expected function, and thus suggesting that FHFs have alternative roles. Indeed, because *FGF13* is expressed in neurons and was associated with intellectual disability (64), an investigation showed how FGF13 stabilizes microtubules in the brain, leading to regulation of growth cones during neuronal development (14). Regulation of microtubules was subsequently described in cardiomyocytes (35, 36). In Das *et al.* we demonstrate that through regulation of microtubule stability FGF13 affects trafficking and targeting of Cx43 connexins, with resultant Cx43 hemichannel-dependent electrophysiological consequences in *cFgf13^KO^*mice. Additional studies have implicated FGF13 in other functions that are independent of VGSCs, including control of neuronal excitability (63) and regulation of the number of caveolae in ventricular cardiomyocytes (43).

Those observations, along with confronting the paradox that the cytoplasmic FHFs affect VGSC SSI (the most commonly reported effect of FHFs on VGSCs, yet a process largely dependent on actions of membrane delimited voltage sensors within the VGSCs), formed the impetus to investigate whether FGF13 affected VGSC function independent of binding. Here, we revealed a novel and unexpected membrane-dependent mechanism controlled by the cytoplasmic FGF13, specifically showing that FGF13 regulated changes in local membrane cholesterol, thereby affecting VGSC SSI. Manipulating membrane cholesterol content alters membrane stiffness and thereby influences voltage-gated ion channel behavior (41, 42). Along with previous observations, our data here showing that *Fgf13* knockout affects local membrane cholesterol in cardiomyocytes, that addition or depletion of cholesterol affects VGSC SSI, and that addition of cholesterol can restore SSI V_1/2_ values in *Fgf13* knockout cardiomyocytes towards WT values, suggests that at least some of the FHF effects on VSGC SSI are mediated by regulation of membrane cholesterol via a VGSC binding-independent mechanism. Cholesterol within the membrane can interact with proteins at Cholesterol Recognition/interaction Amino acid Consensus (CRAC) and inverted CRAC sequences (CARC) (65). Inspection of the cardiac VGSC Na_V_1.5 sequence identifies a CRAC sequence and three CARC sequences within the channel’s transmembrane domains. The CRAC sequence and one of the CARC sequences are in D_III_S_2_ (**Sup Fig 8**), and D_III_ has been implicated in regulating channel inactivation (66). We also examined a cryoEM structure of the homologous Na_V_1.6 that was generated in complex with FGF13 (67). This shows cholesterol molecules adjacent to D_II_ and D_III_ (**Sup Fig 8B**). While there may be differences among specific VGSCs regarding cholesterol interaction, the consistent effects on channel inactivation for those channels studied (40–42, 45) and their sequence homology within their membranes spanning segments suggests that cholesterol interactions and consequent effects are likely similar among the VGSC family.

In contrast to the voltage dependence of SSI, the kinetics of Na_V_1.5 fast inactivation depends upon direct interaction of FGF13 as the binding incompetent FGF13^R/A^ cannot restore the time constant to WT levels when expressed in a c*Fgf13^KO^* cardiomyocyte. Thus, FGF13 appears to affect VGSC kinetics in cardiomyocytes via binding-dependent and independent actions. Because the affinities of FHFs for the Na_V_1.5 cytoplasmic C-terminus are high(30, 57), we posit that an FHF bound to a Na_V_1.5 C-terminus is not a member of the pool that regulates local membrane cholesterol content and thereby affects SSI. In support, our accompanying manuscript identifies a stoichiometric excess of FGF13 in mouse ventricular cardiomyocytes, thus suggesting that a separate pool of FGF13 affects local membrane cholesterol.

Not only do our data support and extend the previous observation (43) that FGF13 affects local membrane cholesterol, but our imaging data with ALOD4 demonstrate that FGF13 is responsible for a heretofore unreported concentration of accessible cholesterol at the IDs compared to other cardiomyocyte membrane areas. The mechanisms for this polarized distribution of accessible cholesterol in cardiomyocytes is not known. Nevertheless, this direct visualization of cholesterol concentrated at the ID is consistent with a previous study in which acute depletion with MβCD specifically disrupted ID integrity within intact hearts (68).

What are the roles of this FGF13-dependent concentration of accessible cholesterol at the ID? We postulate several consequences. First, FGF13 via increasing local accessible cholesterol at the ID supports increased Na_V_1.5 channel availability within the subcellular compartment where Na_V_1.5 channels are found to be concentrated, thus promoting cardiomyocyte excitability, and thereby increased conduction velocity. Indeed, in previous *Fgf13* knockdown (19) or knockout (25, 26, 55) models cardiac conduction was impaired; in our accompanying manuscript we show that *Fgf13* ablation impairs Cx43 targeting, reduces gap junctions at the ID, and contributes to conduction block. Here, we show that loss of FGF13-dependent regulation of local cholesterol at the ID may be another contributor to slowed conduction velocity. Second, we find that the previously reported protection by FGF13 of VGSCs in cardiomyocytes at elevated temperatures (26) is independent of FGF13 binding to Na_V_1.5 and that addition of cholesterol restored the protection to VGSC currents that was absent in c*Fgf13^KO^* cardiomyocytes. Intriguingly, the human *Fgf13* homolog, *FGF12*, was identified as a candidate locus for BrS (20), an arrhythmia exacerbated by febrile illnesses. Perhaps *FGF12* variants associated with BrS offer less protection to Na_V_1.5 currents at elevated temperatures.

Since the FGF13-dependent effects on local membrane cholesterol and on Na_V_1.5 SSI are independent of the interaction with Na_V_1.5 channels, these observations imply that the consequences of FGF13 on local membrane cholesterol are not limited to VGSCs. In a previous study, assessment of the effects of *Fgf13* knockout on ventricular cardiomyocyte action potentials suggested that FGF13 also regulates various cardiac K^+^ channels, yet there was no evidence that FGF13 interacted with specific K^+^ channels (25), consistent with our previous unbiased screen for FGF13 interactors by immunoprecipitation in which we did not identify K^+^ channels (43). Thus, the observed consequences on K^+^ channel currents in the *Fgf13* knockout model likely result from indirect FGF13 effects. As local membrane cholesterol is a well described regulator of cardiomyocyte K^+^ channel currents (69, 70), these observations suggest that the mechanism by which FGF13 affects Na_V_1.5 channels may be more broadly applicable.

Finally, these results suggest that assessment of regulatory effects of ion channel auxiliary subunits should be investigated to determine if all effects depend upon direct binding between the auxiliary subunit and the pore-forming subunit. Many proteins that act as channel auxiliary subunits have identified roles beyond channel regulation. Only separation of function experiments, as we performed with FGF13^R/A^ here, can determine if observed effects on channel function depend upon the direct interaction between the auxiliary subunit and the pore-forming subunit or upon other functions of the auxiliary subunit independent of its binding to the channel pore.

## Methods

### Sex as a biological variable

Since *Fgf13* is an X-linked gene and complete *Fgf13* knockout is lethal (63, 71, 72), the c*Fgf13^KO^* mice studied here were males, obtained as described below.

### FGF13 constitutive knockout animals

This study was approved by the Weill Cornell Institutional Animal Care and Use Committee (protocol no. 2016-0042), and all animals were handled in accordance with the NIH *Guide for the Care and Use of Laboratory Animals*. All mice were maintained on a C57BL/6J genetic background (000664; The Jackson Laboratory). To generate cardiac-specific constitutive knockout mice, we crossed female *Fgf13^fl/fl^*(25) with hemizygous male *Myh6-Cre* (αMyh6-Cre; 011038; The Jackson Laboratory) mice, which have an alpha myosin-heavy chain promoter driving expression of Cre recombinase. Experiments were performed 6-to 12-week-old mice.

### HEK cell transfection for electrophysiology

HEK293 cells were maintained in Dulbecco’s Modified Eagle Medium (DMEM) supplemented with 10% fetal bovine serum (FBS) and 1% penicillin-streptomycin (P/S) under standard culture conditions (37°C, 5% CO₂). For transfection, Lipofectamine 2000 was used according to the manufacturer’s protocol. Cells were transfected with 4 µg of Nav1.5 plasmid along with either 1 µg of GFP or 1 µg of FGF13-GFP or FGF13^R/A^-GFP plasmid. Transfected cells were maintained for 48 hours before being utilized for electrophysiological recordings or immunocytochemical analysis. Data generated from the same batch of cells, transfected simultaneously, were used to create graphs and perform statistical analyses to account for potential batch-to-batch variations.

### Immunoprecipitations of Na_V_1.5 with FGF13 and FGF13^R/A^

HEK293 cells were transfected with 2 µg total of plasmid expressing Na_V_1.5 (30) and 0.4 µg total of APEX-FLAG-V5 tagged FGF13 (VY splice variant) WT or R120A mutant DNA with 6 µl of Lipofectamine 2000 (Thermo Fisher Scientific) and incubated for 24 hours at 37 °C. Cells were washed 3 times with ice-cold phosphate buffered saline (PBS), harvested, and lysed in 0.3 ml of ice-cold Lysis Buffer (20 mM Tris-HCl pH 8, 150 mM NaCl, 2 mM EDTA, 1% IGEPAL® CA-630 (MilliporeSigma)) supplemented with cOmplete™ Mini EDTA-free Protease Inhibitor Cocktail (Roche) and 0.2 mM Phenylmethanesulfonyl fluoride (PMSF). Lysates were centrifuged for 15 min at 17,000 x g at 4 °C. Protein concentration was measured by Bradford assay (Pierce). For the immunoprecipitation, lysates were pre-cleared with 20 µl of pre-equilibrated Protein G magnetic beads (Thermo Fisher Scientific) for 1 h at 4 °C. 50 µl were collected for Western Blot analysis. Pre-cleared lysates were incubated with 2 µl (∼2 µg) of anti V5-Tag monoclonal antibody (Thermo Fisher Scientific, #R960-25) for 16 hours at 4 °C. Immunoprecipitation was performed using 40 µl of pre-equilibrated Protein G magnetic beads (Thermo Fisher Scientific) for 1 h at room temperature. Beads were washed 3 times in lysis buffer and proteins were eluted in 80 µl of Elution Buffer (200 mM Glycine, 50 mM Tris-HCl pH 2.6) immediately neutralized in 40 µl of 1 M Tris-HCl pH 8. Samples were analyzed by SDS-PAGE on a 8-16% Tris-Glycine gel transferred to a PVDF membrane using iBlot3 (Thermo Fisher Scientific). Membranes were blocked in blocking buffer (3% bovine serum albumin [BSA] (m/v), 0.1% TWEEN20 (v/v)) for 1 h at room temperature and incubated overnight with primary antibodies diluted in blocking buffer at 4 °C: Rabbit anti-pan Na_V_ (Alomone ASC-003) 1:1000; Mouse anti-V5-Tag monoclonal antibody (Thermo Fisher Scientific, #R960-25) 1:5000; Mouse anti-GAPDH (Thermo Fisher Scientific, # MA5-15738) 1:1000. Membranes were washed 5 times in TBS washing solution (150 mM NaCl, 20 mM Tris pH 7.6, 0.1% TWEEN20 (v/v)), incubated with secondary antibodies diluted in blocking buffer for 1 h at room temperature: Goat Anti-rabbit IgG, HRP-linked (Cell Signaling, #7074) 1:5000; anti-mouse m-IgGκ BP-HRP (SantaCruz, sc-516102). Membranes were washed 5 times in TBS washing solution, incubated in SuperSignal West Pico PLUS substrate solution (Thermo Fisher Scientifc) and imaged using a ChemiDoc Imager (Bio-Rad).

### Cardiomyocyte Isolation, Cell Culture and Adenoviral Transduction

Ventricular cardiomyocytes were isolated from left and right ventricles *of Myh6-Cre*^-^ (WT) and *cFgf13^KO^* mice using a previously validated Langendorff-free cell isolation method (73). Briefly, after anesthesia with Avertin, the chest was incised to expose the heart and the inferior vena cava, and descending aorta were cut. The heart was then flushed by injecting 10 mL of room temperature “EDTA buffer” in the right ventricle. “EDTA buffer” contains (in mM): 130 NaCl, 5 KCl, 0.5 NaH_2_PO_4_, 10 HEPES, 10 Glucose, 10 2,3-butanedione 2-monoxime (BDM), 10 Taurine, 5 EDTA. After placing an aortic clamp, the heart was excised and transferred to a 60 mm dish containing EDTA buffer. Next, 10 mL of room temperature EDTA buffer was injected into the apex of the left ventricle and the same aperture was used to inject 5 mL of “perfusion buffer” (warmed to 37°C). “Perfusion buffer” contains (in mM): 130 NaCl, 5 KCl, 0.5 NaH_2_PO_4_, 10 HEPES, 10 Glucose, 10 BDM, 10 Taurine, 1 MgCl_2_. After injection of perfusion butter, “collagenase butter” was serially injected (5 × 10 mL) into the left ventricle. “Collagenase buffer” contains (in mg/ml): 0.5 collagenase II, 0.5 collagenase IV, 0.05 protease XIV. The atria were separated and discarded. The ventricles were pulled into ∼1-mm pieces with forceps and cells were dissociated with gentle trituration for 2 min. Enzymatic activity was inhibited by addition of 3 mL of stop buffer (perfusion buffer with 5% sterile FBS, made fresh), and the cell suspension was passed through a 150 mm filter before cells were allowed to gravity settle for 20 min. The cells were then washed twice in perfusion buffer and allowed to settle by gravity for 10 min each before use in various experiments.

Cell culture and adenoviral infection of mouse ventricular myocytes were performed as described (19, 73). Briefly, isolated cardiomyocytes were plated in pre-coated wells with laminin (5 µg/ml in PBS for 1 h at 37°C before laminin was aspirated and well washed with PBS) 12-well plates using plating media (made in M199 [Thermo Fisher Scientific] supplemented with 5% FBS, 10 BDM, penicillin/streptomycin 1x). Cells were allowed to adhere for 1 h before plating media was removed and replaced with 500 µL culture media (made in M199 supplemented with 0.1% BSA, 10 mM BDM, 1 × chemically defined lipid concentrate [Thermo Fisher Scientific], 1 × insulin, transferrin, selenium [ITS, Sigma], P/S. After 2 h in culture media, a subset of *Fgf13^KO^* cardiomyocyte wells was infected with 1.5 µL of Cre-dependent AAV8-DIO-FGF13 virus [adeno-associated virus serotype 8, double-floxed inverted orientation, encoding the FGF13-VY splice variant (63), which is the most highly expressed variant in mouse heart (19)] (1.6 X 10^8^ infectious unit [ifu]/mL). Another subset of KO cardiomyocyte wells was infected with 1.5 µL of AAV8-DIO-FGF13^R/A^ (6.6 X 10^7^ ifu/mL). Cardiomyocytes were incubated in culture media for 48 h for adequate viral expression before subsequent analyses.

### Electrophysiology

All experiments were conducted at room temperature unless otherwise specified. Whole-cell voltage-clamp recordings were performed using a HEKA EPC10 amplifier. Patchmaster software was used to acquire data and Fitmaster software was used to analyze the data. Sodium channel recordings shown in Figures 1 and 2, as well as recordings in HEK293 cells, were performed with an external solution containing (in mM): 124 NaCl, 5 KCl, 2 CaCl₂, 1 MgCl₂, 20 TEA-Cl, 5 HEPES, and 10 glucose, adjusted to pH 7.4 with NaOH. Borosilicate glass pipettes were filled with an internal solution containing (in mM): 125 CsF, 10 NaCl, 10 HEPES, 15 TEA-Cl, 1.1 EGTA, and 0.5 Na-GTP, adjusted to pH 7.3 with NaOH (8). Pipettes were prepared using a P-97 pipette puller (Sutter Instruments) and had a resistance of 0.8–1.2 MΩ.

For sodium channel recordings presented in the remaining figures, cells were perfused with an external solution containing (in mM): 7.0 NaCl, 133.0 CsCl, 1.8 CaCl₂, 1.2 MgCl₂, 11.0 glucose, 5.0 HEPES, and 0.005 nifedipine, adjusted to pH 7.4 with CsOH. The internal pipette solution contained (in mM): 3.0 NaCl, 133.0 CsCl, 2.0 MgCl₂, 2.0 Na₂ATP, 2.0 TEA-Cl, 10.0 EGTA, and 5.0 HEPES, adjusted to pH 7.3 with CsOH (74). Pipettes had a resistance of 2.5–3.0 MΩ. All the components used for external and internal solutions were obtained from sigma.

For current-voltage (I-V) curves, cells were held at −120 mV and depolarized to a series of test voltages ranging from −90 mV to +55 mV. To generate steady-state inactivation curves, currents were elicited at −20 mV for 20 ms following a 500-ms prepulse to test voltages ranging from −120 mV to +20 mV from a holding potential of −120 mV. Normalized currents at the −20 mV pulse were used to construct the steady-state inactivation curve. The data were fitted using the Boltzmann equation:

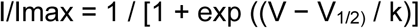

where Imax is the maximum current, V_1/2_ is the half-inactivation voltage, and k is the slope factor. Activation curves were calculated by computing conductance (G, where G=I[(V-Vrev]) from the measured current using a reversal potential (Vrev) calculated for the specific solutions used (see above). The conductance values were then fitted to a Boltzmann equation:

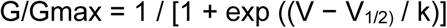

where Gmax is the maximum conductance, V_1/2_ is the half-activation voltage, and k is the slope factor.

To calculate the time constant (τ or tau) of current decay during sodium channel inactivation, the decay phase of the sodium current was fitted to a single-exponential function:

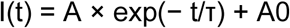

where I(t) is the total current at time t, A is the amplitude of the decaying component of the current at t=0, τ is the time constant of current decay, and A0 is the non-decaying steady-state current. Analyses were performed at different test voltages to determine the voltage dependence of the decay kinetics.

For temperature dependent experiments, the HEK cells and myocytes were initially patched onto at room temperature (RT). After recording the sodium channels current at RT, the temperature of the perfusing bath solution is gradually increased to approximately 37 °C and/or 40 °C and the sodium channel currents were recorded at each temperature levels.

### Cell Imaging

#### Immunocytochemistry with Antibodies

Acutely isolated or cultured cardiac myocytes were fixed with 2% paraformaldehyde for 15 minutes at room temperature. Cells were then permeabilized with 0.2% Triton X-100 in antibody diluent solution for 15 minutes, followed by blocking in antibody diluent solution for 1 hour. Subsequently, cells were incubated with the primary antibody overnight at 4°C. After washing three times with phosphate-buffered saline (PBS), cells were incubated with the secondary antibody for 1 hour at room temperature. Following another three washes with PBS, coverslips containing the stained cells were mounted on slides using mounting medium and allowed to dry before imaging. Antibodies used were rabbit anti-Na_V_1.5 (Alomone, 493-511. 1:500), mouse anti-N-cadherin (Santa Cruz Biotechnology sc-59987, 1:500).

#### Staining with ALOD4

For experiments involving ALOD4 staining, acutely isolated myocytes or HEK293 cells were preincubated with 3 µM ALOD4 (kindly provided by Arun Radhakrishnan (UT Southwestern) conjugated to Alexa Fluor 568 for 20 minutes at 37°C in culture. After incubation, cells were washed three times with PBS and subsequently fixed with 2% paraformaldehyde for 15 minutes. Following fixation, cells were washed three more times with PBS and stained with wheat germ agglutinin (WGA) for 45 minutes at room temperature. Cells were then washed again three times with PBS, mounted on slides using mounting medium, and allowed to dry prior to imaging.

#### In-gel imaging for ALOD4

HEK293 cells were transfected with 1 µg total of eGFP or FGF13VY-eGFP WT or R120A mutant DNA with 3ul of Lipofectamine 2000 (Thermo Fisher Scientific) and incubated for 36 hours at 37 °C. Cells were incubated at 37 °C with Cholesterol or MβCD for 1 h, then with ALOD4 as described above. Cells were washed 3 times in ice-cold PBS, harvested and lysed in ice-cold RIPA buffer (50 mM Tris-HCl, 150 mM NaCl, 0.1% Triton X-100 (v/v), 0.5% sodium deoxycholate (v/v), 0.1% SDS) supplemented with cOmplete™ Mini EDTA-free Protease Inhibitor Cocktail (Roche) and 0.2 mM Phenylmethanesulfonyl fluoride (PMSF). Lysates were centrifuged for 15 min at 17,000 x g at 4 °C. Protein concentration was measured by Bradford assay (Pierce). Samples were analyzed by SDS-PAGE on an 8-16% Tris-Glycine gel. ALOD4 was imaged in-gel using the Alexa547 filter of the ChemiDoc Imager (Bio-Rad).

#### Staining with Filipin

For filipin staining, acutely isolated myocytes were fixed with 2% paraformaldehyde for 15 minutes at room temperature. Cells were washed three times with PBS and incubated with 50 µg/mL filipin solution prepared in PBS for 45 minutes at room temperature. After staining, cells were washed three additional times with PBS and mounted on slides using mounting medium.

#### Imaging

Images of antibody and ALOD4 staining were acquired using a Zeiss LSM 800 confocal microscope. Filipin-stained cells were imaged on a Leica Thunder microscope, with excitation at 405 nm.

#### Image analysis

Line scan analysis was performed to quantify the relative intensity distribution along the myocyte membrane. Using the Profile feature in Zeiss Zen 3.10 Lite software, a 25-pixel-wide line was drawn starting 1 μm before the intercalated disc and extending into the lateral membrane of the cell. Intensity values along the line were quantified for all cells across experimental groups.

For Filipin staining, the mean gray value for each cell was calculated across experimental groups using ImageJ software. These values were used to assess and compare fluorescence intensity levels.

### Preparation and Application of Cholesterol and MβCD Solutions

Cholesterol and methyl-β-cyclodextrin (MβCD) were purchased from Sigma-Aldrich and prepared as previously described (41). Culture media used for HEK cells and myocytes served as the diluent. MβCD was prepared by dissolving the appropriate amount of powdered MβCD in the media. Cholesterol was then added to the solution at a 1:10 molar ratio (Cholesterol:MβCD). The mixture was sonicated for 10 minutes and incubated with continuous rotation in a tube overnight at 37°C. The following day, the solution was filtered through a 0.22 µm filter prior to application to the cells. Cells were treated with either 500 µM MβCD solution for 1 hour to deplete cholesterol or 500 µM MβCD+Cholesterol solution to enrich cholesterol.

### Statistical Analysis

Results are presented as mean ± standard error; the statistical significance of differences between groups was assessed using either a two-tailed Student *t* test or one-way analysis of variance and was set at *P*<0.05. Other statistical analyses were performed as previously described (43).

### Study approval

This study was approved by the Weill Cornell Institutional Animal Care and Use Committee (protocol no. 2016-0042), and all animals were handled in accordance with the NIH *Guide for the Care and Use of Laboratory Animals*. All mice were maintained on a C57BL/6J genetic background (000664; The Jackson Laboratory).

## Acknowledgement

ALOD4-Alexa 568 was provided by permission from Arun Radhakrishnan (UTSW). We also appreciate helpful comments from careful reading by Olaf Andersen (WCM).

